# Targeting regulatory T cells with Interleukin-2 treatment in type 1 diabetes: a response-adaptive, non-randomised, open-label trial of repeat doses of Aldesleukin (DILfrequency)

**DOI:** 10.1101/223958

**Authors:** Eleonora Seelig, James Howlett, Linsey Porter, Lucy Truman, James Heywood, Jane Kennet, Emma L Arbon, Katerina Anselmiova, Neil M. Walker, Ravinder Atkar, Marcin L Pekaiski, Ed Rytina, Mark Evans, Linda S. Wicker, John A. Todd, Adrian P. Mander, Simon Bond, Frank Waldron-Lynch

## Abstract

**Background:** Type 1 diabetes (T1D) results from loss of immune regulation leading to the development of autoimmunity to pancreatic beta-cells, involving autoreactive T effector cells (Teffs). Regulatory T cells (Tregs), that prevent autoimmunity, require Interleukin-2 (IL-2) for maintenance of immunosuppressive functions and, alterations in the IL-2 pathway predispose to T1D. Using an adaptive trial design we aimed to determine the optimal regimen of aldesleukin (recombinant human IL-2) to physiologically enhance Tregs while limiting expansion of autoreactive Teffs.

**Methods:** DILfrequency is a single-center, non-randomised, open-label, response-adaptive study of participants aged 18 to 70 years with T1D. The initial learning phase allocated 12 participants to six different predefined dose-frequency regimens. Then, three cohorts of 8 participants were sequentially allocated dose-frequencies, based on repeated interim analyses of all accumulated trial data. The co-primary endpoints were percentage change in Tregs, Teffs and, CD25 (α subunit of the IL-2 receptor) expression by Tregs, from baseline to steady state. Trial registration ISRCTN40319192 and ClinicalTrials.gov (NCT02265809).

**Findings:** 115 participants were assessed between November 17^th^ 2014 and May 22^nd^ 2016, 38 participants were enrolled with 36 completing treatment. The optimal regimen to maintain a steady state increase in Tregs of 30% and CD25 expression of 25% without Teff expansion is 0.26 × 10^6^ IU/m^2^ (95% CI (−0.007 to 0.485)) every 3 days (1.3 to 4.4). Tregs and CD25 were dose-frequency responsive, while Teffs were not. The commonest adverse event was injection site reaction (464/694 events), with a single participant developing transient eosinophilia at the highest dose (0.47 × 10^6^ IU/m^2^).

**Interpretation:** This response-adaptive trial defined a well-tolerated aldesleukin regimen that specifically induces Treg expansion that can now be trialled to treat T1D.

**Funding:** Sir Jules Thorn Trust, Wellcome, JDRF, SNSF, NIHR

## Introduction

Type 1 diabetes (T1D) is a common chronic disease of children and adults, and yet despite a large investment by both the pharmaceutical industry and academic research communities, insulin replacement remains the only treatment^1,2^. Although, insulin is lifesaving by correcting insulinopenia and resolving hyperglycaemia, it does not treat the underlying autoimmune-mediated destruction of the insulin producing beta cells in the pancreatic islets^3^. To improve short-term and long-term clinical outcomes^4^, there is a need to develop treatments that preserve endogenous insulin production.

Clinical trials of several immunosuppressive agents in T1D demonstrated proof of concept by transiently preserving beta cell function, yet others had no effect or led to disease progression possibly due to incorrect dose selection^5,6^. These results, combined with their side effect profiles, their potential use in the pediatric population and, the requirement for prolonged immunosuppression have prevented the entry of these drugs into clinical practice to treat or prevent T1D. The development of a new non-immunosuppressive therapy that targets the underlying autoimmune cause of T1D would represent an advance on the current standard.

Interleukin-2 (IL-2) is a growth factor, produced by CD4^+^ T effector cells (Teffs) that binds with high affinity to the heterotrimeric IL-2 receptor, which includes the α subunit (CD25)^7^. CD25 is preferentially expressed by IL-2 dependent CD4^+^ T regulatory cells (Tregs). Tregs have an absolute requirement for IL-2 to survive and maintain regulation of Teff responses thereby preventing autoimmunity or collateral tissue damage during immune responses^8^. In T1D, genetic studies have found that several of the genes in the IL-2 pathway contribute to susceptibility while gene-phenotype studies have found that lower expression of CD25 on the surface of Teffs and Tregs is correlated with increased risk of disease^9^. Thus, physiological replacement of IL-2 in T1D to mimic the protection afforded by the risk-reducing alleles of the IL-2 pathway genes could restore Treg mediated immune regulation while avoiding immune activation.

High-dose recombinant Interleukin-2 (rIL-2; aldseleukin) treatment protocols were originally developed to treat metastatic renal carcinoma and melanoma by activating and increasing the numbers of Teffs and natural killer cells^10,11^. More recently, proof-of-concept trials of low-dose aldesleukin have shown some clinical efficiency in chronic graft-versus-host disease (cGVHD), hepatitis C, alopecia areata and systemic lupus erythematosus (SLE)^12–15^. In T1D, therapy with low-dose aldesleukin alone has been shown to be safe^16^, though in combination with rapamycin it caused disease progression^17^. All these studies share an initial induction phase of daily or alternate day dosing that is modeled on the oncology protocols that were developed using standard trial methodologies.

In the previously published DILT1D trial, we employed a state-of-the-art adaptive-dose finding design to define two single doses of aldesleukin that increased frequencies of Tregs by 10% and 20% (0.101 × 10^6^ and 0.497 × 10^6^ IU/m^2^)^18^. In addition, we found that Tregs were desensitized for at least 24 hours after drug administration and, that single doses over 0.380 × 10^6^ IU/m^2^ did begin to activate Teffs. Based on these results we hypothesized that daily dosing was not advantageous due to this desensitization to maintain an increase in Treg frequencies within the physiological range in the absence of Teff expansion. We speculated that a favourable interval might be greater than every two days, but probably not as much as seven or fourteen days.

A next step in this experimental medicine program was to bring the doses determined in the single-dose study forward in a novel response-adaptive repeat dose trial to confirm that a regular regimen of aldesleukin administration could achieve sustained, increased Treg responses without increasing Teff frequencies.

## METHODS

### Study Design and participants

This was a single-centre, non-randomised, open-label, response-adaptive trial of repeated doses of aldesleukin in participants with T1D. The trial commenced with a learning phase where 12 participants were allocated doses and frequencies at the extremes of the available combinations. The dose and frequency for the learning phase were informed by the results of the preceding single dose DILT1D trial ^18^. The next three cohorts of 8 participants were planned to form a tripartite confirming phase. After each cohort completed, all data accumulated in the trial was analysed by the trial statisticians. From the interim reports, the dose frequency committee (DFC) provided decisions regarding the choice of dose and frequency to administer to the subsequent cohorts. The decision-making process of allocating treatment was defined prior to the first meeting of the DFC (appendix).

The study team employed an internet recruitment strategy to enrol eligible participants from across the European Union^19^. Treatment and follow up was performed at the National Institute for Health Research/Wellcome Trust Clinical Research Facility, Addenbrooke’s Hospital, Cambridge, UK. Participants were eligible if they were aged 18 to 70 years and had a duration of T1D ≤5 years from diagnosis. Key exclusion criteria were unstable diabetes with recurrent hypoglycaemia, active clinical infection, active autoimmune thyroid disease, history of severe organ dysfunction, malignancy, history or current or past use of immunosuppressive agents (appendix). All participants provided written informed consent.

The study was done in accordance with the guidelines for Good Clinical Practice and the Declaration of Helsinki. Approval was obtained from the Health Research Authority, National Research Ethics Service (14/EE/1057). The trial was registered at the International Standard Randomized Controlled Trial Number Register (ISRCTN40319192) and ClinicalTrials.gov (NCT02265809). The study protocol was published in advance of the completion and final analysis of the trial ^20^.

### Procedures

Participants were enrolled for 6–16 weeks depending on treatment duration. All participants had 12 visits, including a screening visit, 5–10 treatment visits (dependent on dose-frequency allocation) and a follow-up visit at approximately four weeks after the final dose of drug. For the first learning phase doses of 0.09 and 0.47 × 10^6^ IU/m^2^ of aldesleukin were administered to cohort 1 every 2, 5, 10 or 14 days. Doses and frequencies for subsequent three cohorts were determined by the DFC based on results of interim analyses. Aldesleukin (Novartis Pharmaceuticals UK Ltd) was prepared by the clinical trial pharmacy at Addenbrooke’s Hospital and either administered by a nurse or self-administered under observation. Samples of blood were taken immediately before drug administration for immunophenotyping, full blood counts, and inflammatory markers at all visits. The baseline sample (pre-treatment) was drawn just before administration of the first dose on visit 2, with further samples obtained on following visits within one hour before or after the time recorded on visit 2. To characterise the hyperacute effects of aldesleukin blood samples were also drawn at 90 minutes’ post administration of the drug after the first, second and/or tenth dose. For metabolic assays samples where obtained in fasting or non-fasting state depending on cohort and visit number.

The fluorescence-activated cell sorting (FACS) assay was performed on whole blood following good clinical practice at the Department of Clinical Immunology, Addenbrooke’s Hospital, Cambridge UK within four hours of phlebotomy. Full blood counts, clinical chemistry, metabolic measures (HbA1c, C-peptide, insulin, proinsulin, 1,5 anhydroglucitol), thyroid function tests and autoantibodies (anti-islet, anti-GAD, anti-IA2, anti-ZNT8, anti-TPO, anti-TSH receptor antibodies) were measured at the Departments of Haematology, Biochemistry and Immunology, Addenbrooke’s Hospital, Cambridge UK. IL-2 was measured using the MSD S-PLEX Human IL-2 assay (limit of detection 2 fg/ml) at MSD and converted to IU/ml^21^. Safety was assessed at all visits by adverse event reporting, clinical examination, vital signs and review of the study diary containing daily glucose values and insulin use. A single participant with a previous history of psoriasis had two skin punch biopsies of inflammatory lesions, one during the dosing period (visit 10) and one during the follow up visit (visit 12).

### Outcomes

The co-primary endpoints are the percentage change in Tregs, Teffs and CD25 expression on Tregs from baseline to the average of the last three values immediately preceding drug administration (trough values) when steady state has been achieved. Tregs, Teffs and CD25 expression on Tregs were defined within the CD3^+^CD4^+^ FACS gate^20^ (appendix). Secondary predefined clinical endpoints were change in Treg count and subsets, T effector count and subsets, natural killer cell frequency and count, full blood count and differential, IL-2 and hsCRP levels, autoantibodies, metabolic measures and safety. An exploratory endpoint was the effect of aldesleukin on psoriatic plaques.

### Statistical Analysis

Three populations were predefined for the analyses: The safety population includes all participants who received any dose of aldesleukin. The evaluable population includes participants who received all treatments, where co-primary endpoints were measured. The analysis population is a subset of the evaluable population where the three co-primary endpoints were observed at steady state, used for all primary, secondary and exploratory analysis. Steady state was achieved if the trough value(s) of the percentage increase of Tregs, CD25 or Teffs did not have an upward or downward trend at the end of the dosing schedule. The targets for the co-primary endpoints were set by the DFC after the learning phase. For the primary analysis, a multivariate regression model was fitted with the co-primary endpoints as the dependent variables. The relationship between dose and frequency were analysed in candidate models with the best model having the smallest Akaike information criterion (AIC).

The calculated dose/frequency to achieve the target increases in Treg and CD25, was obtained by setting the linear predictors for Treg and CD25 to equal the targets and solving the simultaneous equations for dose and frequency. The precision to which dose can be administered is 0.01×10^6^ IU/m^2^ and practically frequency can only be integer days meaning that the predicted calculated dose/frequency to achieve the increases is unlikely to be practical. Therefore, the joint probability that all 3 co-primary endpoints fell within the target ranges was calculated for each dose/frequency able to be practically administered. In addition to the joint probability, the Mahalanobis distance, a measure of how close each dose/frequency is to achieving the targets in Treg and CD25 was calculated. The practical dose/frequency that maximised the joint probability was selected as the optimal dose/frequency, if the Teffs at this administration were predicted to be within the target range. The optimal dose/frequency will be reported in the results.

For the secondary endpoints, two measures were defined, the change and the percentage change. The change is the difference between the baseline and time point measurements. The percentage change is defined as the ratio of the change to the baseline measurement. For the 90-minute variable, a linear mixed effects model was fitted with the outcome defined as the difference between the measurement and the corresponding trough value with dose, frequency and dosing visit included as covariates in the model as well as a random intercept with participant as the grouping variable. For the other derived variables, a univariate regression model was fitted including the same covariates as the primary analysis model. As the effects of dose and frequency cannot be separated, the R^2^ value and a global F-test will be reported to evaluate how much of the variability in the endpoint is explained by dose and frequency. Where the secondary outcome measures were not measured repeatedly at each visit the same measures were defined at visit 12. Continuous variables are expressed as mean (±SD, N) with the SD only reported when N≥4 and discrete variables as count (%).

### Role of the Funding Sources

Funding was provided by the Sir Jules Thorn Trust, the Swiss National Science Foundation (SNSF), Wellcome, JDRF, MRC Biostatistics Unit and, NIHR Cambridge Biomedical Research Centre. The funding bodies had no role in the study design, data collection and analysis, decision to publish, or preparation of the manuscript. The corresponding author (FWL) had full access to all the data, carried out the analysis of the primary and secondary outcomes with JH and SB (trial statisticians) and had the final responsibility for the decision to submit to publication.

### Results

Between November 17^th^ 2014 and May 22^nd^ 2016, 38 participants were enrolled at the NIHR Wellcome Trust Clinical Research Facility, Addenbrooke’s Hospital, Cambridge, UK (figure 1). The baseline characteristics of the safety population (N=37) are presented in table 1. When cohort 1 (12 participants) completed the first learning phase, a second learning phase was conducted, as not enough participants in the trial had achieved steady state for the primary endpoints. Cohort 2 (8 participants) was treated with two doses of 0.09 and 0.47 × 10^6^ IU/m^2^ aldesleukin at frequencies of between 3 and 10 days (figure 2A). The DFC determined that steady state had been achieved in enough participants to set a Treg target of a 30% increase (range ±5%), a CD25 target of a 25% increase (range ±5%) and a Teff target of a 0% increase/decrease (range −100%, ∞) from baseline for remainder of the trial. For the confirming phase of the trial, doses of 0.2 or 0.32 × 10^6^ IU/m^2^ every 3 days were allocated to cohort 3 (8 participants) (figure 2B), with a dose of 0.32 × 10^6^ IU/m^2^ every 3 days to the final cohort (8 participants) (figure 2C).

**Figure 1.**
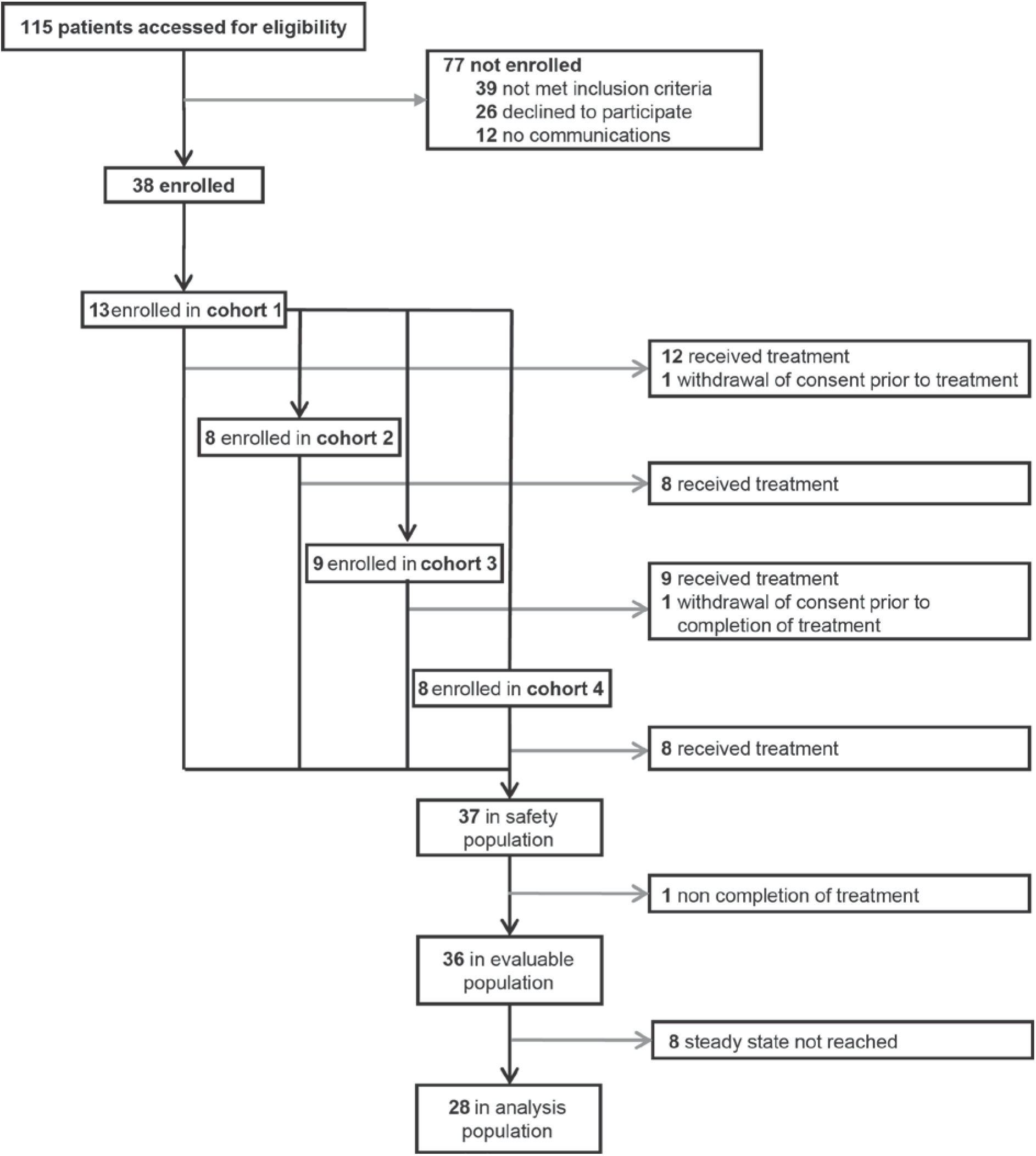
Trial profile

**Figure 2.**
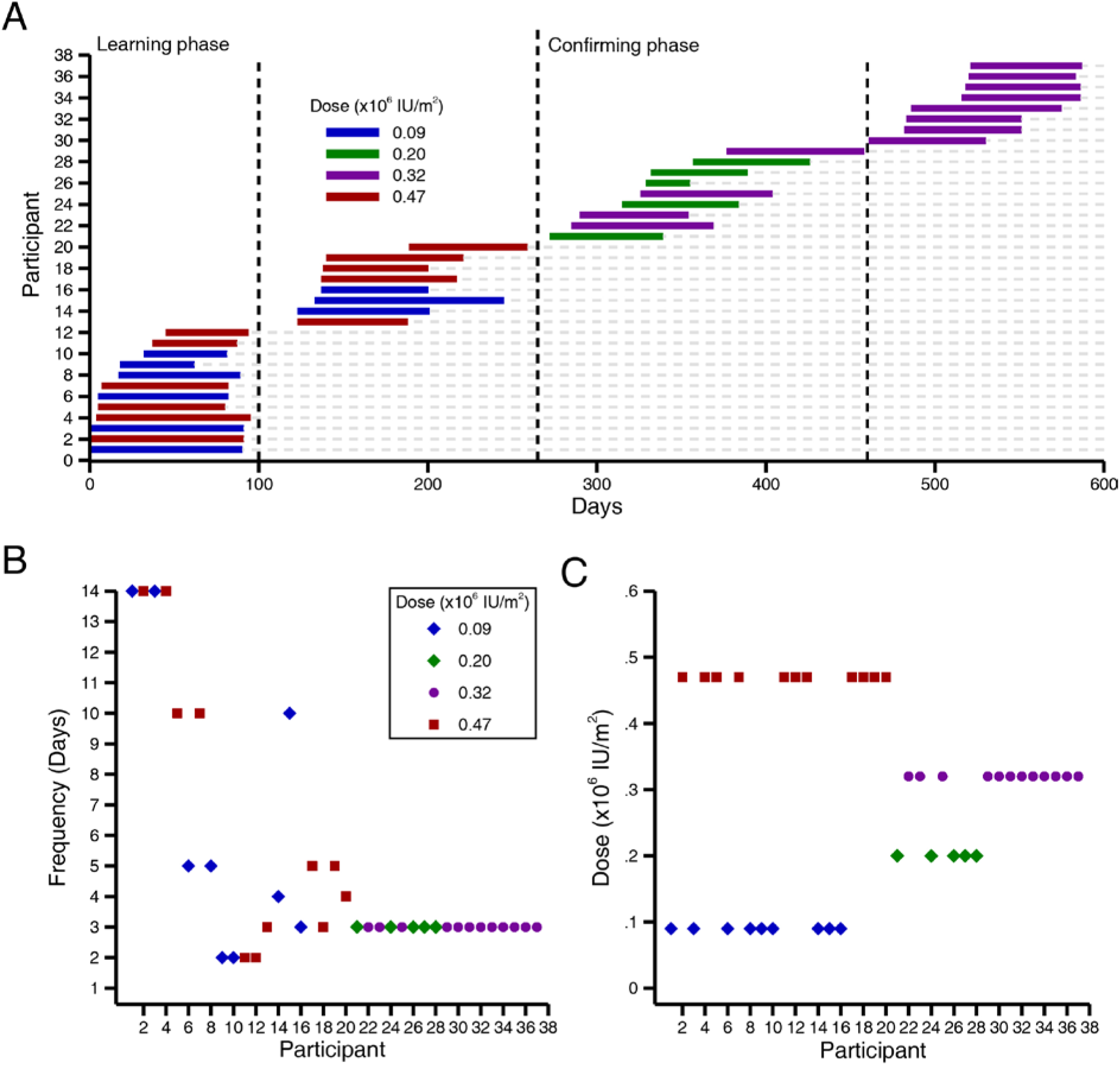
DILfrequency adaptive design, dose and frequency allocation. (A) The study was conducted in two phases, a learning phase (259 days) and confirming phase (315 days), in the first learning phase participants were allocated either 0.47 × 10^6^ or 0.09 × 10^6^ IU/m^2^ doses of aldesleukin with sequential allocations of the longest dose interval first. After the first 12 participants, an interim analysis was performed that determined the dose and frequency allocation for the next cohort of 8 participants. The process was then repeated after completion of each cohort with dose-frequency allocations based on analysis of all data collected from previous cohorts. (B) Participant frequency allocation showing convergence of the trial to every 3-day administration. (C) Participant dose allocation showing convergence of the trial to the 0.02 and 0.32 × 10^6^ IU/m^2^ dose to estimate the optimal dose.

**Table 1.** Baseline characteristics of safety population (N= 37)

| Characteristic |  |  |
| --- | --- | --- |
| Age (years) | Mean (SD) | 38.2 (±11.2) |
|  | Median (min-max) | 36 (19-65) |
| Age group | 18-35 years | 15 (40.5%) |
|  | 36-65 years | 22 (59.5%) |
| Sex | Male | 22 (59.5%) |
|  | Female | 15 (40.5%) |
| Diabetes duration (months) | Mean (SD) | 20.85 (±13.70) |
|  | Median (min-max) | 21.1 (1.3-59.1) |
| Diabetes duration | ≤100 days | 1 (2.7%) |
| | >100 days and ≤2 years | 20 (54.1%) |
| | > 2 years and ≤5 years | 16 (43.2%) |
| Age at diagnosis (years) | Mean (SD) | 36.5 (±11.3) |
|  | Median (min-max) | 35 (17-61) |
| BMI (kg/m<sup>2</sup>) | Mean (SD) | 23.878 (±3.369) |
|  | Median (min-max) | 23.88 (19.27-33.27) |
| BSA (m<sup>2</sup>) | Mean (SD) | 1.804 (±0.197) |
|  | Median (min-max) | 1.75 (1.47-2.20) |
| Type 1 diabetes-associated Ab | Anti-islet positive | 12 (32.4 %) |
|  | Anti-GAD positive | 31 (83.8 %) |
|  | Anti-IA2 positive | 15 (41.7%) |
|  | Anti-ZnT8 positive | 19 (52.8%) |
| Thyroid associated Ab | Anti-TPO positive | 8 (22.2%) |
|  | Anti-TSH positive | 0 (0 %) |
| Plasma glucose (mmol/l) | Mean (SD) | 9.60 (±3.76) |
|  | Median (min-max) | 9.1 (3.5-18.1) |
| 1,5 anhydroglucitol (ucg/ml) | Mean (SD) | 6.32 (±5.50) |
|  | Median (min-max) | 4.43 (0.96-26.15) |
| HbA1c (mmol/mol) | Mean (SD) | 59.4 (±17.5) |
|  | Median (min-max) | 56 (34-100) |
| HbA1c (%) | Mean (SD) | 7.584 (±1.603) |
|  | Median (min-max) | 7.27 (5.26-11.30) |
| Insulin (pmol/l) | Mean (SD) | 284.6889<br>(±544.4443) |
|  | Median (min-max) | 93.353 (0.000-3032.323) |
| Proinsulin (pmol/l) | Mean (SD) | 3.0769 (±3.8757) |
|  | Median (min-max) | 2.049 (0.000-16.012) |
| C-peptide (pmol/l) | Mean (SD) | 290.8 (±239.3) |
|  | Median (min-max) | 225 (0-949) |
| Daily insulin dose (Units/kg body weight) | Mean (SD) | 0.425 (±0.240) |
|  | Median (min-max) | 0.38 (0-1.15) |

There were no serious adverse events (AE) reported in the trial, with 37 (100%) of participants reporting an AE (table 2). All the AE’s were mild (670/694, 96.5%) or moderate (24/694, 3.5%) with the most common being hypoglycaemia, injection site nodule and injection site erythema (table 2; appendix). A single participant on dose 0.47 × 10^6^ IU/m^2^ every 4 days developed an asymptomatic eosinophilia on visit 7 that resolved by visit 11. Pre-existing eosinophilia improved by visit 11 or even resolved by visit 10 in two participants receiving 0.32 × 10^6^ IU/m^2^ every 3 days, and 0.09 × 10^6^ IU/m^2^ every 2 days, respectively (appendix). In some participants (21/37, 56.8%) there were transient decreases in lymphocytes at 90 minutes after the first dose and again after the second (11/20, 55.0%) or tenth dose (11/16, 68.8%) without the development of lymphopenia. There was no evidence of the development of thyroid dysfunction despite enrolling participants with positive anti-TPO antibodies (8/36, 22.2%), with no participants developing de novo anti-TPO or anti-TSH antibodies following treatment (appendix). Aldesleukin had no effect on hsCRP, renal, bone or liver biochemistry and clinical FACS parameters (appendix). One participant with stable psoriasis was treated with 0.32 ×10^6^ IU/m2 every 3 days without any exacerbating of pre-existing disease (appendix). Overall, treatment was well tolerated at all doses and frequencies with no participants discontinuing the study or being withdrawn by the trial team due to AE’s or reactions.

**Table 2.** Adverse events by severity and type in the safety population (N= 37)

| Type of Adverse event | Category | Events | Participants |
| --- | --- | --- | --- |
| Total | Serious adverse events (N) | 0 (0%) | 0 (0%) |
|  | Adverse events | 694 (100%) | 37 (100%) |
|  | AEs events by severity |  |  |
|  | - Mild<br>- Moderate | 670 (96.5%)<br>24 (3.5%) | 37 (100%)<br>13 (35.1%) |
| Related AEs | Related adverse events by severity (N/%) | 474 (99.0%)<br>5 (1.0%) | 35 (94.6%)<br>3 (8.1%) |
|  | - Mild<br>- Moderate |  |  |
|  | All related AEs |  |  |
|  | - Acne | 2 (0.4%) | 2 (5.4%) |
|  | - Asthenic condition | 3 (0.6%) | 3 (8.1%) |
|  | - Dry eye | 1 (0.2%) | 1 (2.7%) |
|  | - Eosinophilia | 1 (0.2%) | 1 (2.7%) |
|  | - Injection site erythema | 230 (48.0%) | 35 (94.6%) |
|  | - Injection site nodule | 234 (48.9%) | 33 (89.2%) |
|  | - Malaise | 1 (0.2%) | 1 (2.7%) |
| Related AEs | - Migraine headaches | 1 (0.2%) | 1 (2.7%) |
|  | - Nasopharyngitis | 3 (0.6%) | 3 (8.1%) |
|  | - Nausea and vomiting symptoms | 2 (0.4%) | 2 (5.4%) |
|  | - Oropharyngeal pain | 1 (0.2%) | 1 (2.7%) |
|  | Related adverse events (N/%) by |  |  |
|  | - Expected | 472 (98.5%) | 35 (94.6%) |
|  | - Unexpected | 7 (1.5%) | 5 (13.5%) |
|  | Related adverse events by cohort |  |  |
|  | Cohort 1 |  |  |
|  | - Mild | 112 (100%) | 11 (91.7%) |
| Related AEs | - Moderate | 0 (0%) | 0 (0%) |
|  | Cohort 2 |  |  |
|  | - Mild | 100 (98%) | 7 (87.5%) |
|  | - Moderate | 2 (2%) | 1 (12.5%) |
|  | Cohort 3 |  |  |
|  | - Mild | 121 (98.4%) | 9 (100%) |
|  | - Moderate | 2 (1.6%) | 1 (11.1%) |
|  | Cohort 4 |  |  |
|  | - Mild | 141 (99.3%) | 8 (100%) |
|  | - Moderate | 1 (0.7%) | 1 (12.5%) |
| Hypoglycaemia | Events per number of glucose measurements | 94/9159 (1%) |  |
|  | Events per number of participants |  | 20 (54.1%) |

In the analysis population, the model including the effect of dose, dosage frequency, dosage frequency squared, and the dose-frequency interaction fitted the data best with the smallest combined AIC of 696.92. The model explained 50.35% of the variability in Tregs (p=0.0022), 76.12% of the variability in CD25 (p<0.0001), and 37.32% of the variability in Teffs (p=0.0246). The optimal dose of aldesleukin is 0.26 × 10^6^ IU/m^2^ (95% CI −0.007, 0.485) every 3 days (95% CI 1.3, 4.4). The probability of this treatment regime achieving the targets for the 3 endpoints is 0.742 or 74.2% (Figure 3: appendix).

**Figure 3.**
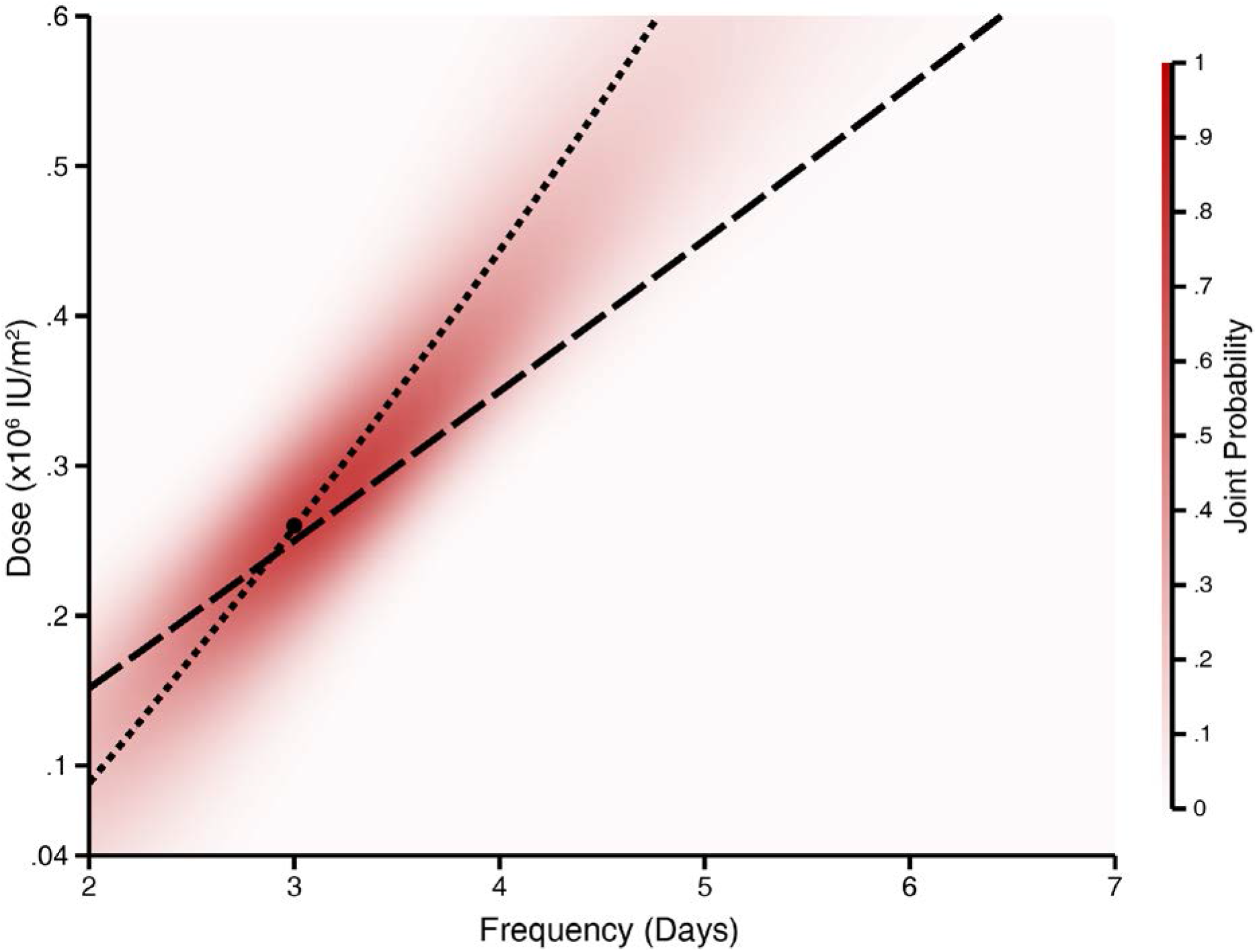
Primary Endpoints. This statistical model described the Treg and Treg CD25 dose-frequency response best, with the larger dashed line showing the dose-frequencies predicted to achieve increases from baseline of 30% for Tregs and smaller dashed line showing a 25% increase for Treg CD25. The dot marks the optimal dose-frequency with the joint probability of achieving shown as a red density plot.

Analysis of the 3-day dosing interval found that the 0.47 and 0.32 × 10^6^ IU/m^2^ doses produced similar mean Treg increases of 41.31% (N=2) and 45.87% (±22.43, N=8) respectively at the primary endpoint when administered every 3 days, while the 0.20 × 10^6^ IU/m^2^ dose increased Tregs 20.34% (±11.05, N=4) (figure 4A-C). A maximum increase in Tregs of 83.67% from baseline was observed on visit 9 after 8 doses of 0.32 × 10^6^ IU/m^2^ every 3 days. This increase was maintained up to visit 11 to give an overall mean increase of 79.59% for the primary endpoint in this participant (figure 4B, C). The lowest dose, 0.09 × 10^6^ IU/m^2^ was only effective in increasing Tregs when administered every 2-days; wider dosing intervals with this dose had no effect (figure 4A-C; appendix Treg FACS figures). At 5-day interval only the highest dose of 0.47 × 10^6^ IU/m^2^ was effective in maintaining a Treg increase of 23.28% (N=2) becoming a no-effect dose at 10 or 14-days (figure 4C).

**Figure 4.**
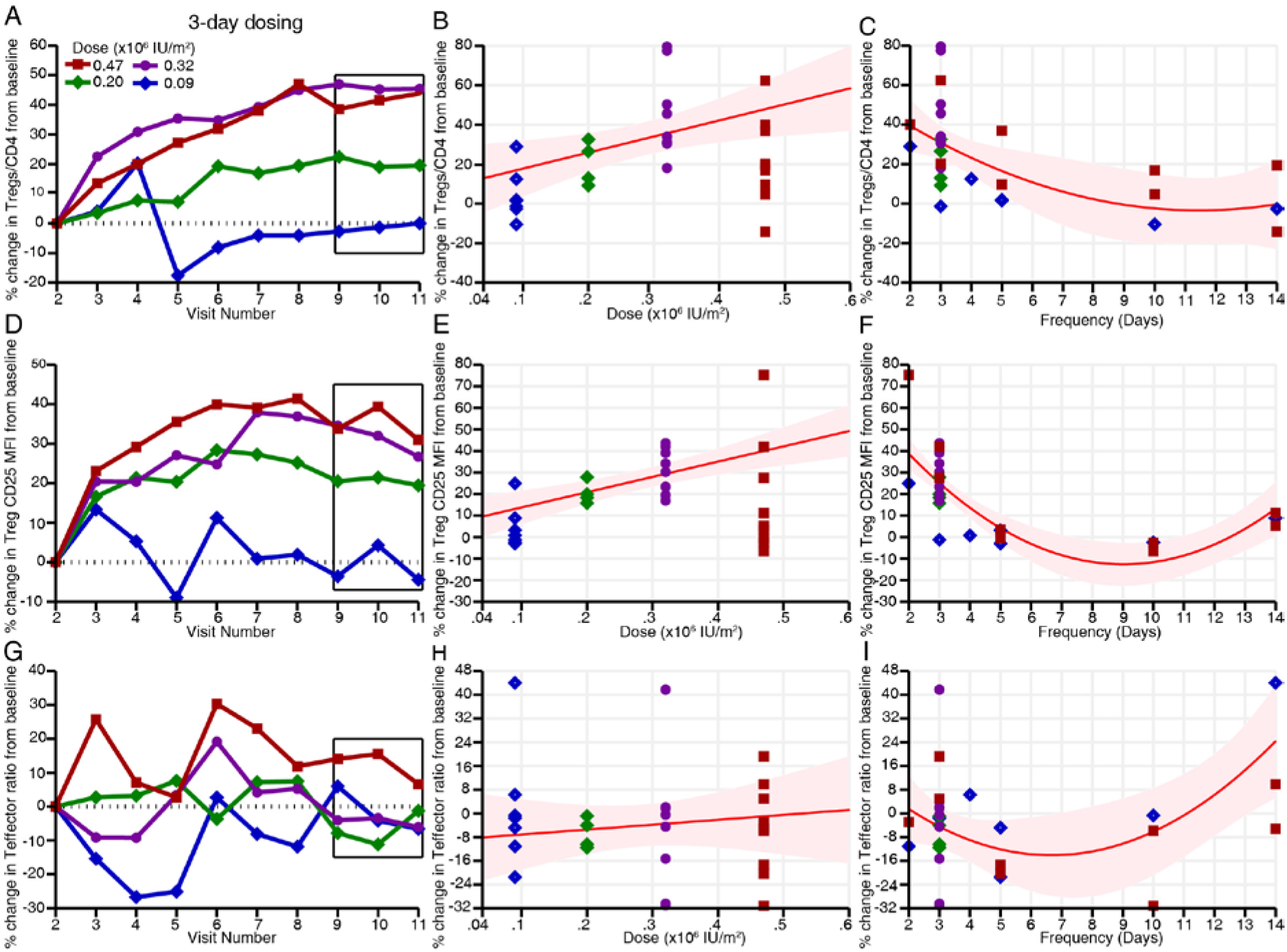
Regulatory T cell, Regulatory T cell CD25 and T effector responses to aldesleukin treatment. (A) Regulatory T cell (Treg) proportions as a percentage of CD4^+^ T cells for doses administered every 3 days with the box highlighting the final three trough values at steady state that were used to define the Treg primary endpoint (average response plots shown across the four doses) (B) The percentage change in Tregs at the primary endpoint for all doses administered with the predicted dose response at the best frequency (every 3 days, red line) showing an increase in precision of the estimates around the optimal dose (0.26 × 10^6^ IU/m^2^, shaded area 95% CI). (C) The increase in Tregs for all frequencies allocated with the predicted frequency response at the optimal dose (0.26 × 10^6^ IU/m^2^, red line) showing increased precision of the estimates around day 3. (D) Increase in CD25 expression on Tregs at the 3-day frequency at the primary endpoint. (E and F) The change in Treg CD25 at all doses and frequencies with increased precision of the estimates around the 3-day frequency (red line). (G) The changes in the T effector (Teff) ratio of proportion naïve effectors to memory effector at the 3-day frequency. (H and I) The change in Teff ratio at the primary endpoint at all doses and frequency, showing no dose-frequency response at 0.26 × 10^6^ IU/m^2^ every 3 days.

The expression of CD25 increased on Tregs with the maximal response by dose 6.1 (±2.3, N=15) at 3-day dose intervals (figure 4D-F; appendix). At the 3-day interval the lowest dose failed to maintain an increase of Treg CD25 expression at steady state, while the 0.20, 0.32, and 0.47 × 10^6^/m^2^ achieved a sustained increase of CD25 expression of 20.46 (±5.20, N=4), 31.09 (±10.32, N=8), and 34.72% (N=2) respectively (figure 4F). Increase of the dose interval to greater than 3 days resulted in loss of the Treg CD25 response at the time points assessed (figures 4F). The changes in Treg proportion and CD25 expression from baseline were dependent on both dose and frequency (R^2^=0.054, p<0.0001). There was no clear effect of dose on Teffs at 3-day dosing intervals with doses of 0.09, 0.20, 0.32, 0.47 × 10^6^/m^2^ achieving percentage increases from baseline of −1.58 (N=1), −6.74 (±5.11, N=4), −4.51 (±23.08, N=8), and 12.07% (N=2) respectively (figure 4G, H, I).

Prior to treatment, IL-2 levels in participants (N=36) were 0.0025 (±0.0019) IU/ml (41.82 (±30.91) fg/ml) comparable to those previously reported for healthy individuals and T1D patients^18^. The maximal sustained increase in IL-2 was at the 0.47 × 10^6^/m^2^ dose delivered every 2 days with a mean increase from baseline to plateau (trough values before last 3 doses) of 0.0247 IU/ml (appendix). At the 3-day interval dosing, participants administered the 0.20 and 0.32 × 10^6^ IU/m^2^ doses had mean IL-2 increases at plateau of 0.0003 (±0.0003, N=4) and 0.0029 (±0.0038, N=8) IU/ml, respectively (figure 5A). A single participant had an isolated peak of IL-2 to 2.95 IU/ml on visit 6 that corresponded on review with an episode of gastroenteritis that was recorded as an AE (figure 5A), this increase in endogenous IL-2 is similar to what we have previous measured in acute self-limiting viral gastroenteritis^22^. The overall effect of the repeated doses of 0.20 and 0.32 × 10^6^ IU/m^2^ aldesleukin every 3 days was to increase IL-2 levels by 18.75% (±21.38, N=4) and 162.23% (±165.57, N=8) above baseline at plateau. At 5-day interval only, the 0.47 × 10^6^ IU/m^2^ dose produced an increase of 0.0005 IU/ml (N=2), when measured 5 days after each dose, while there was no increase with lower doses or at 10- or 14-day intervals (figure 5B, C). IL-2 levels observed at the measured time points are consistent with the single dose levels observed in our previous study^18^ with no evidence of drug accumulation or unexpected drug elimination due to the induction of receptor-mediated clearance.

**Figure 5.**
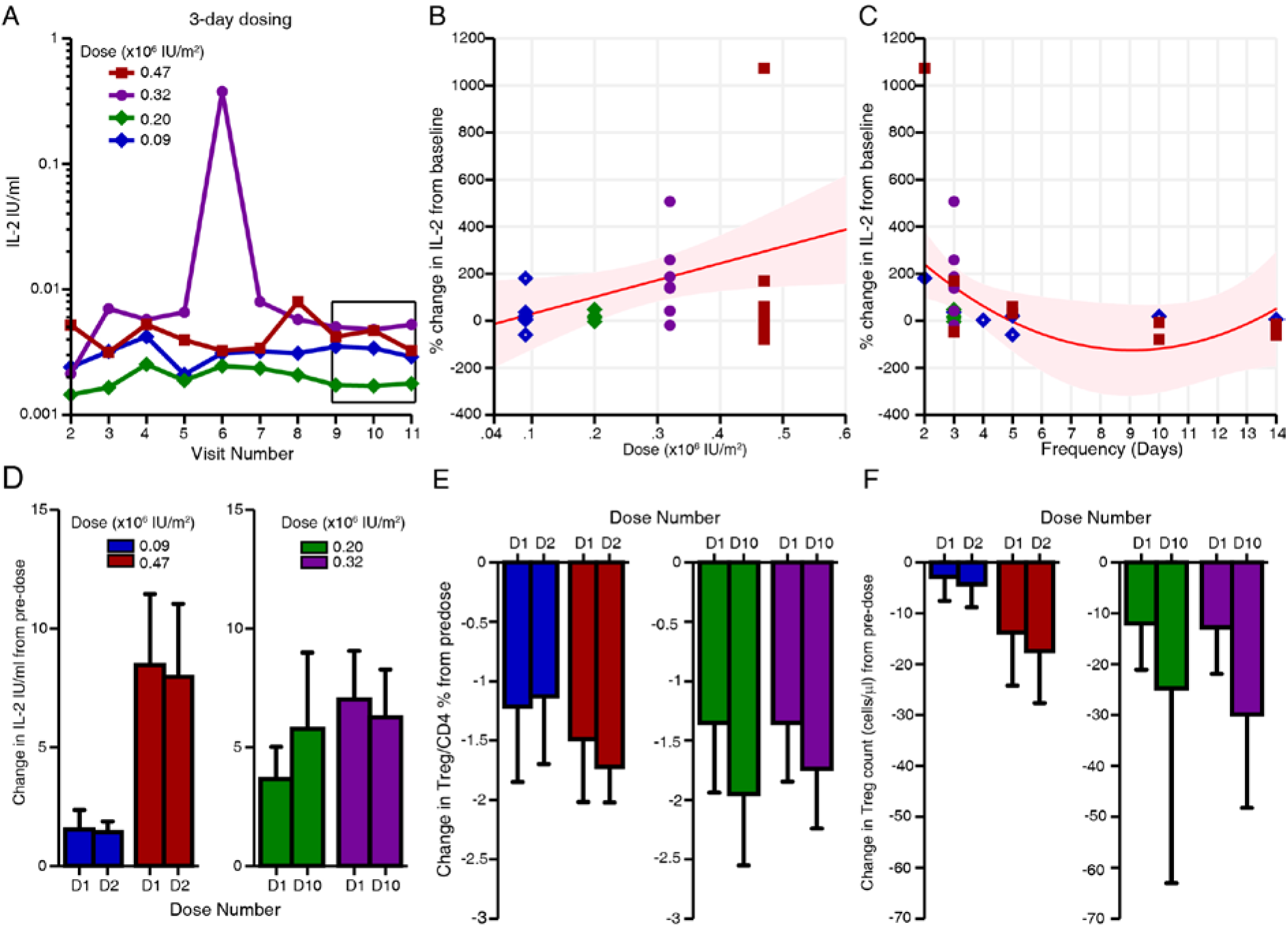
Interleukin-2 levels and hyperacute Regulatory T cell responses. (A) Interleukin (IL-2) levels during treatment with aldesleukin every 3-days with the box highlighting the final three trough values. The peak on visit 6 in the 0.32 × 10^6^ dose group is coincident with the report of an adverse event (gastroenteritis) from a participant (average response plots shown across the fours doses). (B and C) The percentage change in IL-2 levels from baseline and at the measurement of the primary endpoint for all doses, with the estimated dose response at the best frequency (every three days, red line) and the optimal dose (0.26 × 10^6^ IU/m^2^, red line, shaded area 95% CI). (D) Change in IL-2 levels at 90 minutes following the first dose (D1), the second dose (D2) or the tenth dose (D10). (E) The decline in regulatory T cells (Treg) proportions as a percentage of CD4^+^ T cells at 90 minutes. (F) The decline in Treg counts at 90 minutes at the commencement of treatment and when Tregs have increased by dose 10 (visit 11) (mean and bars showing 95% CI).

At 90 minutes following the first dose, there was a dose dependent increase in IL-2 levels that ranged from 0.7487 to 13.9685 IU/ml (N=28, dose coefficient=18.25, p<0.001), consistent with those observed in the previous single dose study (DILT1D)^18^. There was no difference in the levels of IL-2 at 90 minutes after the second dose of either 0.09 and 0.47 × 10^6^ IU/m^2^ with mean increase of 1.43 (±0.48, N=7) and 7.97 IU/ml (±4.00, N=9) respectively (figure 5D). To assess if the increase of Tregs and CD25 expression at steady state would alter IL-2 levels, they were measured 90 minutes after the tenth doses of 0.2 and 0.32 × 10^6^ IU/m^2^ on visit 11, with mean increases 5.77 (±2.02, N=4) and 6.26 IU/ml (±2.41, N=8) respectively (coefficient=0.20, p=0.7001) (figure 5D), thereby establishing that the immune system will be recurrently exposed to transient nonTreg-specific levels (>0.015ml/IU)^18^ after administration. Simultaneously at 90 minutes, there was a rapid transient reduction in the percentage of Tregs in circulation (−1.36%; ±0.60; N=28) after the first dose, this reduction was observed again following the second (−1.46%; ±0.57; N=16) or tenth dose (−1.81%; ±0.53; N=12) when Tregs have reached steady state (Figure 5E). Though the proportion of Tregs trafficking out of the circulation remained similar regardless of the number of doses administered, the reduction in Treg count was greater at steady state (Figure 5F).

The Treg count was 54.9 (±22.6, N=28) ul/ml before treatment with the mean increase from baseline of 17.3 (±7.9, N=4), 30.0 (±13.5, N=8) and 29.5 (N=2) cells/uL at doses of 0.20, 0.32 and 0.47 × 10^6^/m^2^ every 3 days respectively (Figure 6A). The increase in Treg count was only observed when aldesleukin was administered every 2, 3 or 5-days. The changes in Treg counts from baseline to steady state was dose-frequency dependent (r^2^=0.4108, p=0.0130). Within the Treg population both naïve and memory Tregs were increased by 39.60% (±36.45, N=28) and 59.39% (±48.11, N=27), respectively, from baseline to steady state, with the 0.2 and 0.32 ×10^6^ IU/m^2^ doses resulting in the greatest increases of effector memory (EM) Tregs 50.23% (±44.16, N=11) (figure 6D). Within the rest of the CD4 compartment, there was no effect of dose or frequency of administration on total CD4 (r^2^=0.1640, p=0.3680) or CD4 T effector counts (r^2^=0.1582, p=3890). Similarly, there was no change in NK cell count from baseline to steady state (0.028 cell/uL; ±0.072; N=28) with no dose-frequency reponse (r^2^=0.1564, p=0.3957) (appendix).

**Figure 6.**
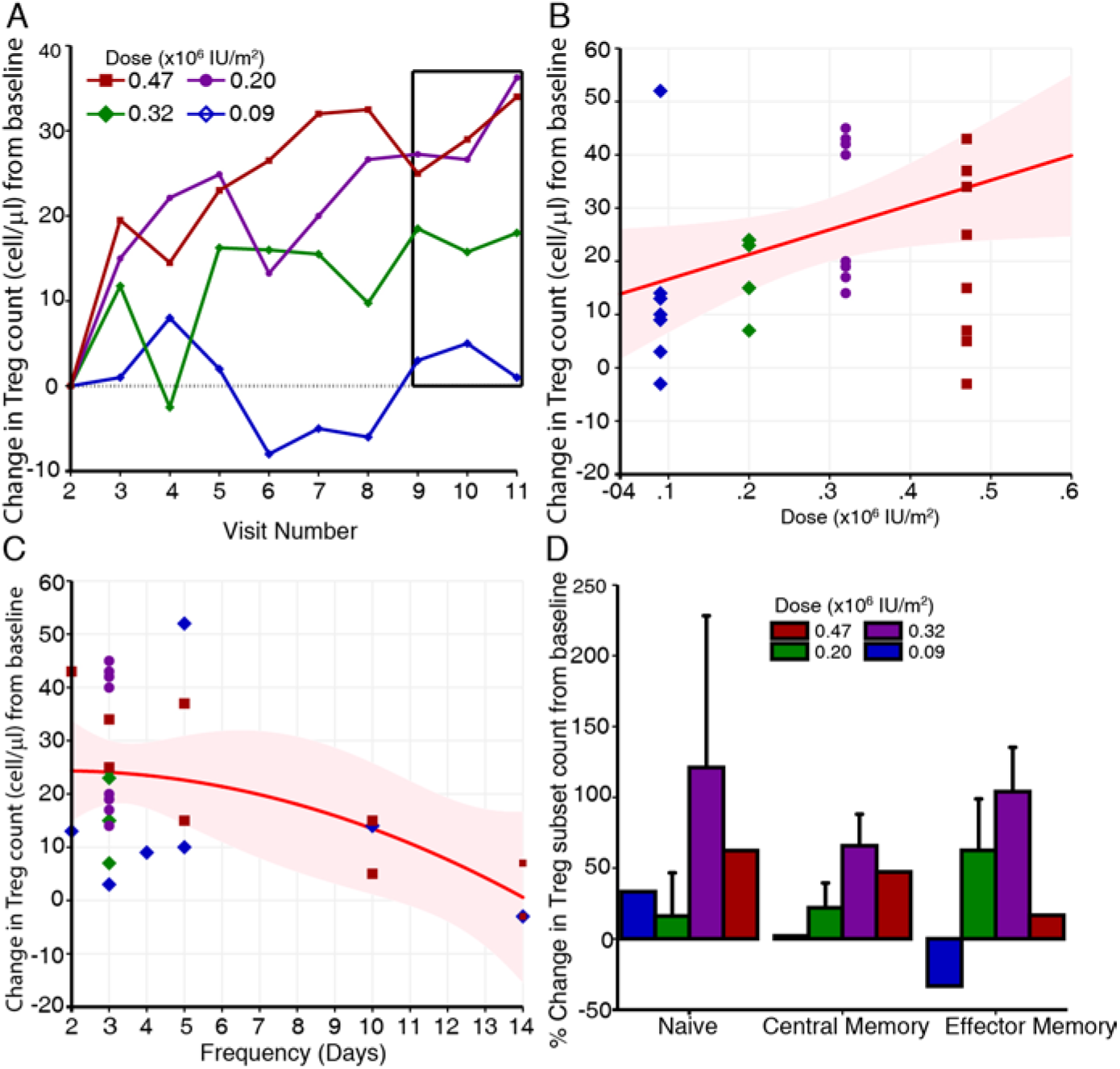
Regulatory T cell counts and subsets. (A) Regulatory T cell counts during treatment with aldesleukin every 3 days with the box highlighting the final three trough values. (B and C) The changes in Treg count at all doses and frequencies allocated showing the estimated dose response when aldesleukin is administered every 3 days (red line) and frequency response at 0.26 × 10^6^ IU/m^2^ (red line, shaded area 95% CI). (D) The change in Treg subsets from baseline to measurement of the primary endpoint (mean and bars showing 95% CI).

Metabolic control remained stable throughout the trial with fasting C-peptide slightly increasing from baseline to the final measurement by 26.75% (±68.42, N=25) with no effect of dose or frequency of aldesleukin administration observed (figure 7A-C). 1,5 Anhydroglucitol, a marker for short term glucose control, proinsulin to C-peptide ratio, a marker for β-cell stress, and random glucose, remained stable with a baseline change of 17.63% (±42.65, N=23), 6.07% (±44.34, N=16), and 7.99% (±36.92, N=26), respectively. Insulin use at baseline and follow-up was 0.421 units/kg (±0.227, N=27) and 0.464 units/kg (±0.193, N=26), respectively (figure 7D-F). Baseline and follow-up HbA1c were 56.1 (±15.5, N=28) and 51.6 mmol/mol (±12.3, N=28) respectively meaning that there was small improvement in blood glucose control (−4.71% (±8.55, N=28) (appendix).

**Figure 7.**
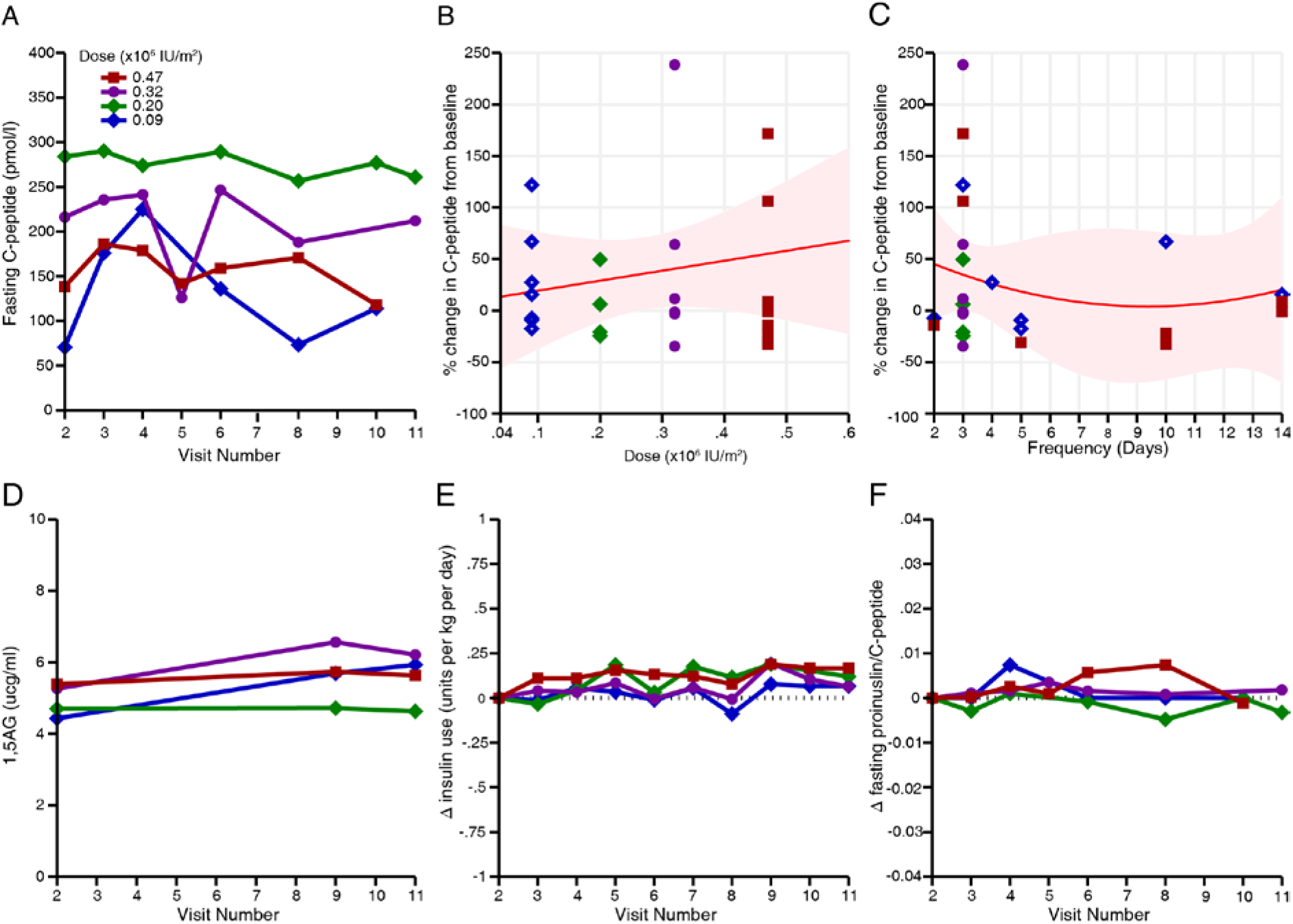
Metabolic parameters. (**A**) Fasting C-peptide levels pre-treatment and during treatment (B and C) Percentage change in fasting and non-fasting C-peptide levels for all allocated dose-frequencies with the predicted dose response at the best frequency (3 days, red line, shaded area 95% CI) and frequency response at the optimal dose (0.26 × 10^6^ IU/m^2^, red line). (D) 1,5-anhydroglucitol level before and during treatment, with increased levels representing better glycaemic control (E and F) insulin use, and fasting proinsulin to C-peptide ratio for all doses administered every 3 days (average response plots shown across the four doses).

### Discussion

The DILfrequency study has established the optimal dose and administration frequency to maintain an increase in Tregs and, Treg CD25 expression while not expanding Teffs in participants with T1D. A response adaptive design has been employed to develop an immunomodulatory treatment protocol that increases immune regulation within physiological levels to restore health while preserving pathogen responses in participants with T1D. This experimental medicine approach of defining a treatment regimen based on our understanding of the immunopathogenisis of the disease prior to testing clinical efficacy contrasts with current strategies of conducting traditional randomised, double-blind placebo control trials that test efficacy first, with limited success^23,24^.

The adaptive-response design of DILfrequency by continuously adapting the dose and frequency after each interim analysis defined the “optimal” dose-frequency to achieve the trial targets. This methodology allowed for the flexibility to adapt to a wider range of dose frequencies than could be pragmatically investigated in a fixed dose study of a similar size. This enabled the development of a regimen where 0.26 × 10^6^ IU/m^2^ of aldesleukin every three days, should result in the targeted expansion of Tregs by 30%, increase of Treg CD25 by 25% without increasing Teffs. Lower doses and/ or lower frequencies induced an attenuated or even no response of Tregs while higher doses and/or higher frequencies risk Treg desensitisation, expansion of Teffs and are less practical for patients. This regimen of aldesleukin differs from other schemes, currently administered in T1D clinical trials, where a 5-day induction treatment (0.5–1 × 10^6^ IU/day for children or 1 × 10^6^ IU/day for adults) is followed by single dose every 7 or 14 days dependent on treatment cohort^25^. Our regimen is similar to a dosing regimen (0.1 to 0.2 × 10^6^ IU/m^2^ three times per week) that was utilised to treat another clinical population, namely paediatric patients after allogeneic hematopoietic stem cell transplantation ^26^

Treatment with these doses and frequencies of aldesleukin used in DILfrequency was well tolerated with most common AE’s being mild injection site reactions and, all infections were self-limiting. Eosinophilia, a well-known side effect of IL-2 treatment, only occurred asymptomatically in a single participant receiving the highest dose repeatedly, while pre-existing eosinophilia in two other participants improved or even resolved. Another side effect of IL-2 treatment, especially in cancer treatment, is the development of autoimmune diseases such as thyroiditis^27^. No secondary autoimmune diseases were triggered even in participants with high pre-existing risks such as TPO-autoantibody positivity. Moreover, we were able to observe in detail how a case of pre-existing psoriasis remained stable throughout the trial^28^. Overall, the aldesleukin treatment regimen had a favourable safety profile, which is essential for any potential therapy in T1D.

In DILfrequency, administration of repeated doses of 0.47 × 10^6^ IU/m^2^ of aldesleukin every two days, increased IL-2 levels to a plateau above the Treg specific level of 0.015 IU/ml and, could activate Teff cells^18^ while the IL-2 concentrations at the 0.2 and 0.32 × 10^6^ IU/m^2^ doses every three days were increased to above pretreatment levels. Previously we observed that IL-2 levels increased to peak concentrations that are not selective for Tregs at 90 minutes and 24 hours after administration of aldesleukin^18^. Here we found that these peak drug concentrations at 90 minutes were replicated at all doses and frequencies, even after several administrations when Tregs had expanded to a steady state This suggests that there is minimal target-mediated drug disposition/clearance induced by the dosing regimens tested.

There was a transient decrease in both Treg percentage and count in the circulation at 90 minutes, consistent with our previous observation following a single dose of aldesleukin where Tregs were decreased at 90 minutes and at 24 hours following dosing. This could indicate increased retention of Tregs in the tissues or vasculature upon aldesleukin administration or recruitment of Tregs from the blood to tissue, or both. Interestingly, once Tregs had expanded in response to the repeated aldesleukin doses, the reduction in the number of Tregs in the circulation at 90 minutes increased though the percentage decrease (~20%) was unchanged from initial visits. This suggests that there is a Treg population that retains its original sensitivity to aldesleukin. Further detailed analyses are required to understand the immunophenotype and function of these Tregs. The measurement of Treg count also enabled a comparison with the responses to IL-2 treatment in cGVHD where there is partial clinical efficacy^29^. In cGVHD, the pre-treatment Treg count was approximately threefold lower than that observed in T1D and increased, after 12 weeks of treatment (1 × 10^6^ IU/m^2^/daily) to a Treg plateau within the range achieved in DILfrequency with three-day dosing. These more intensive aldesleukin regimes may be required for the autoinflammatory orders such as cGVHD and SLE where there is a reduction Treg count as compared to T1D where Treg function alone is impaired ^12,15^.

DILfrequency was designed to develop an optimal IL-2 treatment regimen for T1D based on a primary immunological outcome rather than metabolic efficiency. The informativeness of the metabolic endpoints measured are limited owing to the short period of treatment but did show that there was no evidence of adverse alterations of these metabolic parameters or evidence of beta cell stress. There was a small increase of endogenous fasting C-peptide^30^, stable glucose homeostasis with no change in insulin usage by participants in the trial. The analysis of the trial was also limited to peripheral blood from adult T1D patients due to ethical and practical considerations.

In DILfrequency, an adaptive design has been successfully employed to develop a well-tolerated IL-2 intermittent dosing regimen of 0.26 × 10^6^ IU/m^2^ of aldesleukin every three days that targets and maintains homeostatic Treg responses without inducing short-term immunosuppression. The clinical efficacy of this self-administered treatment can now be evaluated in a substantially larger trial using a longer duration of the optimised dosing regimen in T1D patients with residual insulin production, to determine if these immunomodulatory effects will induce remission from beta cell autoimmunity, leading to preservation of endogenous insulin production to improve clinical outcomes.

### Contributions

FWL served as the chief investigator and wrote the first draft of the manuscript, other members of the writing team included ES, JH, AM and SB. All authors where involved in the conduct of the study and collection of the data. JH, AM and SB performed the study analysis. All authors participated review in the article and approved the final version.

### Conflicts of Interest

FWL has received fees for consulting and speaking on type 1 diabetes and immunotherapeutics from Epidarex Capital, GlaxoSmithKine, Novo Nordisk, Eli Lilly and Hoffmann-La Roche. The CamT1D team that FWL leads has received research and clinical trial funding from Hoffmann-La Roche and GlaxoSmithKine. JAT has received advisory board fees from Pfizer, Celgene and AstraZeneca.

## Acknowledgements

The study was sponsored by the University of Cambridge and Cambridge University Hospital Trust. The authors acknowledge the support of the National Institute for Health Research (NIHR) Cambridge Biomedical Research Centre and the Cambridge Clinical Trial Unit (CCTU) trial coordination; the NIHR/Wellcome Trust Clinical Research Facility, Addenbrooke’s Centre for Clinical Investigation (ACCI) for clinical facilities; Department of Clinical Immunology, Addenbrooke’s Hospital, Cambridge, UK; The JDRF/Wellcome Trust Diabetes and Inflammation laboratory sample processing team led by Helen Stevens, information technology, administration team led by Judy Brown and data teams at the Cambridge Institute for Medical Research; Dr Laurence Peterson, JDRF/Wellcome Diabetes and Inflammation laboratory, Cambridge Institute for Medical Research; Dr. Kevin O'Shaughnessy, Department of Medicine, University of Cambridge, independent chair of the DILfrequency trial steering committee. For assistance with identification of potential participants we acknowledge the ADDRESS-2 study and colleagues at the Wolfson Diabetes and Endocrine clinic, Addenbrooke’s Hospital, Cambridge, UK. The authors thank Prof Ken Smith, Department of Medicine, University of Cambridge, for the critical review of the manuscript. The generous contribution of the participants in DILfrequency is very gratefully acknowledged.

## Data Availability Statement

The data cannot be anonymised sufficiently to be able to be put in the public domain without the risk of participant identification. Data is available on request through the Cambridge University institutional repository <u>www.repository.cam.ac.uk/handle/1810/265634</u>.

